# Universal base editing for hemophilia B

**DOI:** 10.1101/2024.11.13.623331

**Authors:** Nemekhbayar Baatartsogt, Yuji Kashiwakura, Takafumi Hiramoto, Rina Ito, Rikako Sato, Yasumitsu Nagao, Hina Naruoka, Haruka Takata, Morisada Hayakawa, Khishigjargal Batjargal, Tomoki Togashi, Atsushi Hoshino, Taro Shimizu, Yusuke Sato, Tatsuhiro Ishida, Osamu Nureki, Tsukasa Ohmori

## Abstract

The repair of pathological gene variants is an ultimate aim for treating genetic diseases; however, it is not practical to develop different therapeutic reagents for each of the many variants that can occur in a gene. Here, we investigated whether base editing to induce a gain-of-function variant in blood coagulation factor IX (FIX) can increase FIX activity as a treatment strategy for hemophilia B. We engineered a G:C to A:T substitution at c.1151 of *F9* by cytosine base editing to generate R338Q, known as the Shanghai *F9* variant, which markedly potentiates coagulation factor activity. An adeno-associated virus vector harboring the base editor converted more than 60% of the target G:C to A:T and increased FIX activity in HEK293 cells harboring patient-derived *F9* variants, as well as in knock-in mice harboring a human *F9* cDNA. Furthermore, administration of lipid nanoparticles embedded with the base editor mRNA and gRNA increased FIX activity in mice. These data indicate that cytosine base editing to generate R338Q in FIX can become a universal genome editing strategy for hemophilia B.

## Main text

Genome editing therapy using CRISPR-Cas9 has much potential as an innovative treatment for genetic diseases. Cas9 nuclease recognizes DNA bound to guide RNA (gRNA), and catalyzes a double-strand break (DSB) in the target gene^1^.DSBs induce non-homologous end joining, a main repair mechanism of DSBs that can cause nucleotide deletion or insertion, resulting in gene knockout^2^. Clinical trials of genome-editing therapies that use this technique for transthyretin amyloidosis and hereditary angioedema have been performed, and have obtained favorable therapeutic effects^3,4^. Furthermore, genome editing therapy targeting *BCL11A* in hematopoietic stem cells for sickle cell disease and thalassemia has recently been approved by the UK Medicines and Healthcare Products Regulatory Agency and the US Food and Drug Administration^4,5^. However, genome editing accompanied by DSBs has the potential to be genome toxic through the generation of large deletions and chromosomal translocations and rearrangements^6^. Therefore, base editing technologies that do not involve DSBs are being developed, such as base editing, epigenome editing, and prime editing.

Base editing is a technology that can repair pathological single-nucleotide variants. It involves Cas9 nickase conjugated with a deaminase to enable C:G to T:A or A:T to G:C substitution^7^. Fifty-five percent of all pathological variants in human genetic diseases are single nucleotide variants, and 60.6% of these are A:T to G:C and C:G to T:A variants^8^. Therefore, approximately 33% of all genetic diseases can potentially be treated by base editing^9^. Repair of pathological variants by base editing is a goal of personalized therapy, but it is not practical to prepare individual reagents for each variant. Therefore, base editing therapies that can cover many patients by editing a single base are now being proposed. For example, approaches to repair a common variant in sickle cell disease to the non-pathological Makassar β-globin^10^ and to disrupt the *PCSK9* splice acceptor to treat familial hyperlipidemia^11^ have proceeded toward clinical application.

Hemophilia is a congenital bleeding disorder caused by variation in the blood coagulation factor VIII (*F8*) or IX (FIX) (*F9*) genes. The incidence of hemophilia was estimated at 1 in 5,000 males^12^, and the World Federation of Hemophilia estimated a worldwide hemophilia patient population of 400,000^13^. Compared with hemophilia A (abnormality of *F8*), hemophilia B (abnormality of *F9*) involves a higher proportion of single-nucleotide variants^14^, making it a particularly promising target for base editing. We have previously repaired a single nucleotide variant in *F9* in induced pluripotent stem cells derived from hemophilia B patients^15^. To apply base editing technology to many hemophilia B patients, we hypothesized that engineering a gain-of-function variant in *F9* through base editing would be effective. In this study, we present a technique to increase FIX activity by *in vivo* base editing that will enable the treatment of many hemophilia patients.

## Results

### FIX R338Q substitution increases the activity of hemophilia B FIX variants

The success of adeno-associated virus (AAV) vector-mediated gene therapy for hemophilia B is attributed to exploitation of the FIX Padua variant (R338L). FIX Padua was discovered in familial thrombophilia patients and exhibits eight times higher FIX activity than wild-type FIX^16^. Although R338 is conserved in mammalian evolution, amino acid substitution at R338 increases FIX activity^17^. To increase FIX activity by *in vivo* base editing, we focused on substitution at R338. As shown in Fig. 1a, we obtained the R338Q amino acid substitution by a G:C to A:T transition at c.1151 of *F9*. This variation is known as FIX Shanghai, which causes inherited thrombosis^18^. We first compared activity among wild-type, R338Q, and R338L FIX in HEK293 cells transduced with pcDNA3 expressing each *F9* cDNA. We found that R338Q and R338L have 4- and 8-fold increased FIX activity compared with wild-type FIX, respectively (Fig. 1b). AlfaFold prediction of binding between coagulation factor VIII (FVIII) and activated FIX (FIXa) showed that R338Q and R338L FIX variants created novel noncovalent binding to the A2 domain of FVIII (Extended Data Fig. 1). This is consistent with the enhanced coagulation activity of FIX Padua requiring the cofactor activity of FVIII^19^. We then constructed cDNAs of FIX variants associated with mild and moderate types of hemophilia B (Extended Data Table 1) without or with R338Q and compared FIX activity in HEK293 cells. We observed increased FIX activity in the cell supernatant with the introduction of R338Q in a variety of variants (Fig. 1c and 1d). These data indicate that the introduction of R338Q by base editing can increase FIX activity in many hemophilia B patients.

**Fig. 1.**
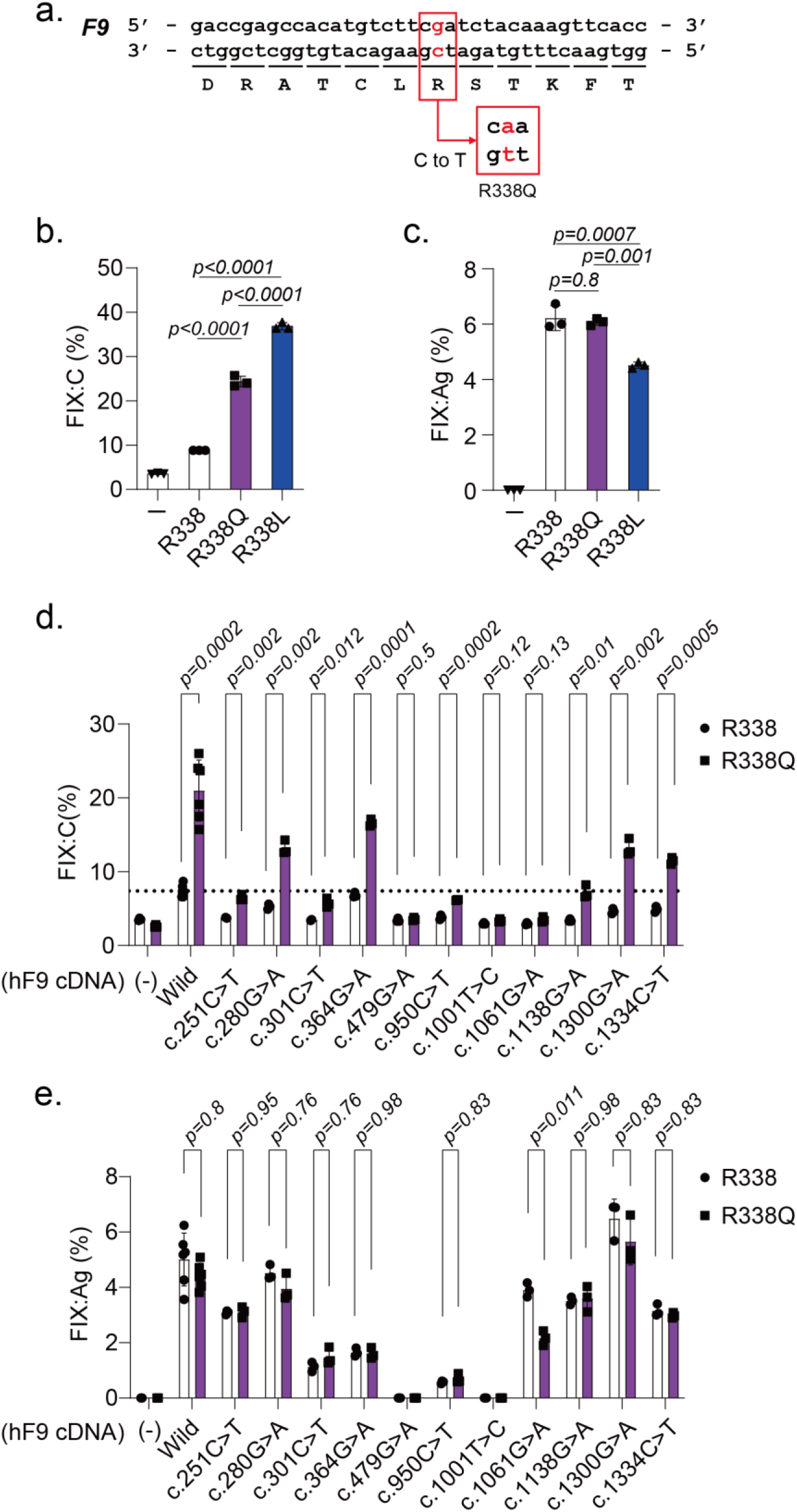
R338Q substitution increases FIX activity. **(a)** Nucleotide and amino acid sequences around the target site. C>T conversion of the minus strand at c.1151 of *F9* generates R338Q. **(b, c)** FIX activity (FIX:C) **(b)** and FIX antigen (FIX:Ag) **(c)** in the supernatant of HEK293 cells transfected with pBApo-EF1α expressing the indicated FIX cDNA [mean ± SD (n = 3)]. Statistical analysis was conducted by one-way ANOVA followed by Tukey’s *post hoc* test for pairwise comparisons. **(d, e)** Increase in FIX:C in hemophilia B variants by the R338Q substitution. The FIX:C **(d)** and FIX:Ag **(e)** in the supernatant of HEK293 cells transfected with pBApo-EF1α expressing the indicated FIX cDNA without or with R338Q substitution [mean ± SD (n = 3–6)]. Statistical analysis was conducted using *t*-tests.

### Base editor for engineering FIX R338Q

There was an NGG and an NG sequence on the 3′ side of the target site of SpCas9 and SpCas9-NG, respectively^20^. These corresponded to the PAM sequence (Fig. 2a). We considered the target site suitable for C to T conversion by apolipoprotein B mRNA editing enzyme (APOBEC)-based C to T base editing. The editing window was typically 4–8 nucleotides numbered from the 5′ end of the gRNA^9^. To determine the optimal gRNA sequence for the introduction of R338Q, we compared DSB efficiency at the target site using two gRNA sequences and SpCas9 or SpCas9-NG in HEK293 cells. gRNA1 combined with SpCas9 was the most efficient combination for inducing DSBs (Fig. 2b). Off target analysis by Guide-seq showed that SpCas9 had fewer off-target sites compared with SpCas9-NG (Fig. 2c). We therefore selected gRNA1 and SpCas9 for further analysis. Of note, the off-target sites did not contain any C nucleotides within the target range of APOBEC (Fig 2a).

**Fig. 2.**
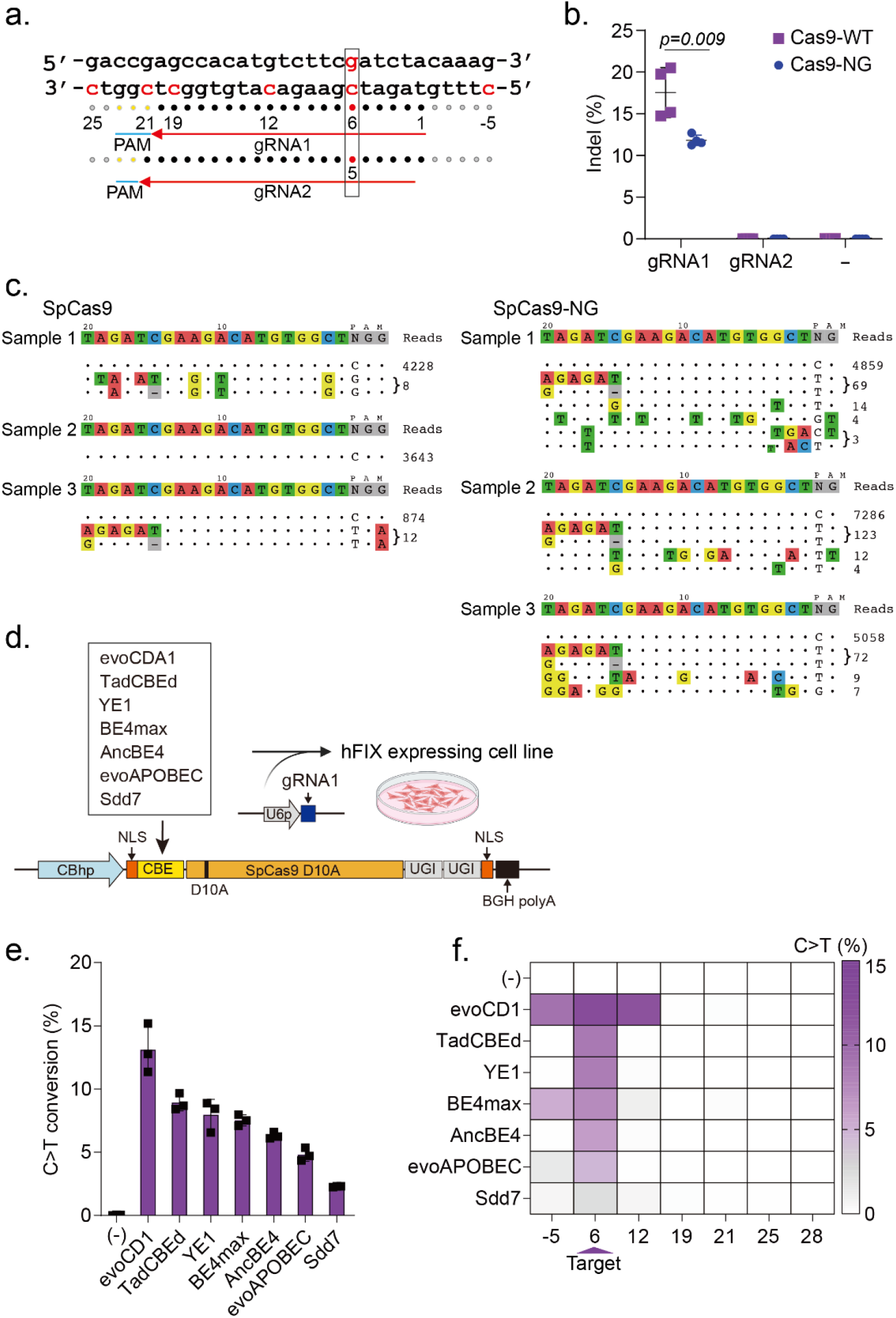
Selection of gRNA sequence and base editor. **(a)** gRNA sequences and the target site for the introduction of R338Q in human *F9*. **(b)** HEK293 cells were transfected with pX330 (SpCas9 or SpCas9-NG) and a plasmid expressing the indicated gRNA. Quantitative analysis of indel frequency assessed by T7 endonuclease assays [mean ± SD (n = 4)]. Statistical analysis was conducted using *t*-tests. HEK293 cells were transfected with pX330 (SpCas9 or SpCas9-NG) containing a gRNA1 expression cassette together with dsODN. The off-target site was analyzed by Guide-seq. The results from three samples are shown. **(d, e, f)** HEK293 cells stably expressing wild-type human *F9* were transfected with pX330 conjugated with a base editor and a plasmid expressing gRNA1. **(d)** Schematic diagram of the experiment. **(e)** Induction of C>T editing at the mRNA target site assessed by NGS [mean ± SD (n = 3)]. **(f)** Visualization of mean bystander C>T conversion.

We next tried to select a suitable base editor for the introduction of R338Q into *F9*. We constructed plasmids expressing SpCas9 (D10A) conjugated with G:C to A:T base editors, including evoCDA1, YE1, BE4max, AncBE4, evoAPOBEC^9^, TadCBEd^21^, and sdd7^22^. HEK293 cells expressing a human *F9* cDNA were transfected with the plasmids, and then G:C to A:T transition efficiency in *F9* mRNA was assessed by next generation sequencing (NGS) (Fig. 2d). evoCDA1 showed the most efficient base editing, but also had the highest bystander effect (Fig. 2e and 2f).

Genome wide off-target effects caused by base editors can be observed at sites unrelated to the gRNA, especially in transcribed regions that display single-stranded DNA within the R-loop^23^. To assess gRNA-independent off-target effects of base editors, we employed an R-loop assay, which generates R-loop structures by the expression of catalytically dead SaCas9 with gRNAs targeting unrelated sites. We transduced HEK293 cells with plasmids harboring dead SaCas9 (D10A and H840A) and gRNA sequence (site 3, 4, 5, and 6), and SpCas9 (D10A) conjugated with a base editor (TadCBEd, BE4max, or evoCDA1) and gRNA1 for R338Q induction. TadCBEd showed less bystander C>T conversion around gRNA1 for R338Q induction (Extended Data Fig. 2). We also detected marginal gRNA-independent off-target base editing with TadCBEd, compared with BE4max and evoCDA1 (Extended Data Fig. 2).

We next constructed a dual AAV vector system to express SpCas9 (D10A) conjugated with BE4max or TadCBEd by protein splicing with an intein (Fig. 3a). Transduction of HEK293 cells expressing *F9* cDNA with an AAV6 vector harboring an indicated base editor succeeded in editing the target site in a concentration-dependent manner (Fig 3b). The base editing efficacy at the target site by TadCBEd was significantly higher than that by BE4max (Fig 3b). BE4max exhibited some bystander editing around the target site (Fig 3b). Unexpected base edits other than G:C to A:T were less than 10% (Fig. 2c).

**Fig. 3.**
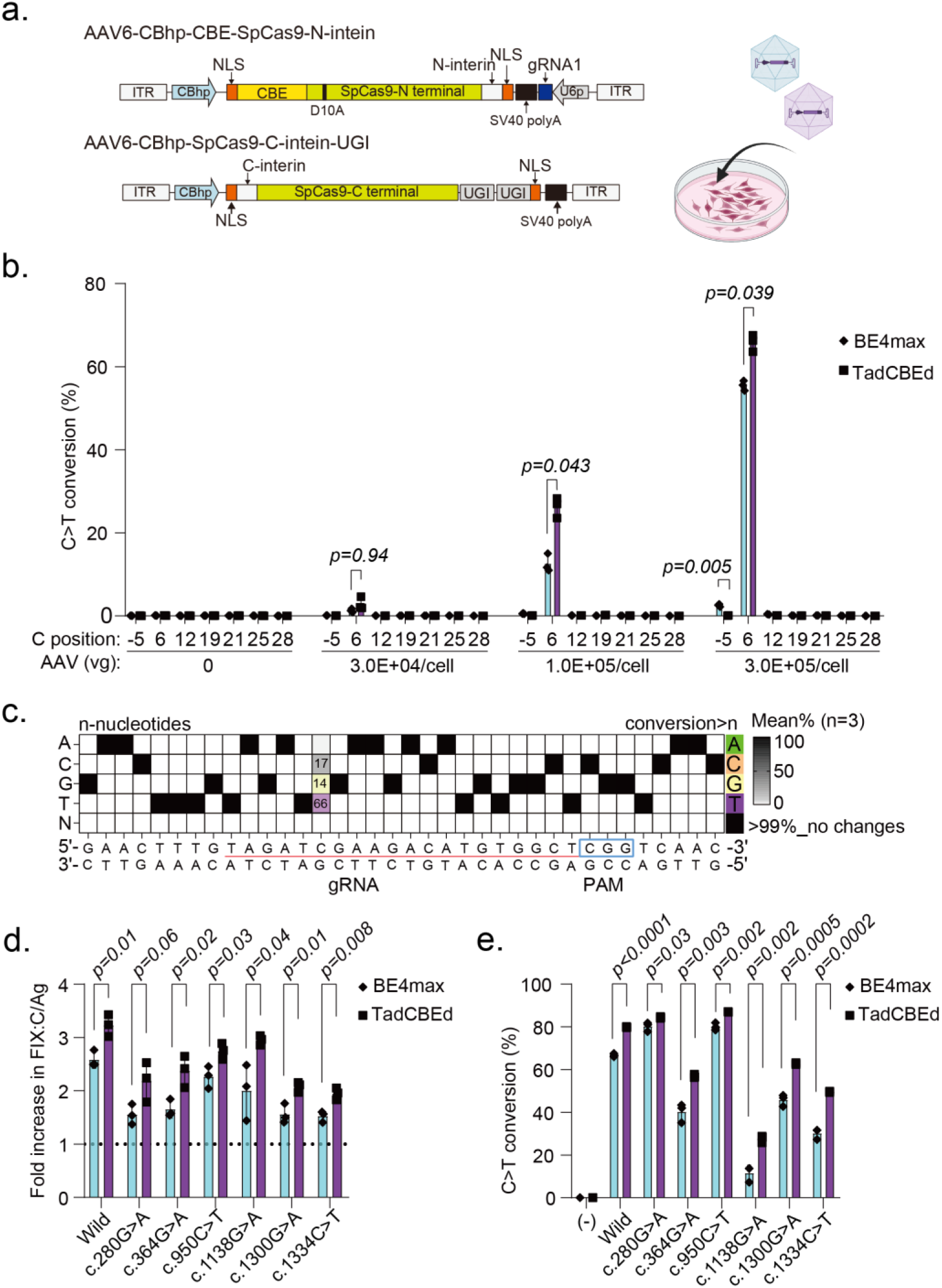
Base editing *in vitro* increases FIX activity. HEK293 cells stably expressing a human *F9* cDNA were transduced with two AAV6 vector systems that express either SpCas9 (D10A) conjugated with TadCBEd base editor or BE4 max and UGI sequences, together with the gRNA1 sequence. **(a)** Schematic diagram of the experiment. **(b)** Induction of C>T editing of mRNA around the target site assessed by next generation sequencing in HEK293 cells expressing a wild-type human *F9* cDNA [mean ± SD (n = 3)]. Statistical analysis was conducted using *t*-tests. **(c)** The mean conversion frequency other than C>T substitution. **(d)** Increase in FIX activity (FIX:C) and antigen (FIX:Ag) ratios following the transduction of HEK293 cells expressing a wild-type human *F9* cDNA with hemophilia B variant *F9* cDNAs ([mean ± SD (n = 3)]. Statistical analysis was conducted using *t*-tests. **(e)** Induction of C>T editing of mRNA around the target site assessed by next generation sequencing [mean ± SD (n = 3)]. Statistical analysis was conducted using *t*-tests.

We then assessed the base editing efficacy and the increase in FIX activity in HEK293 cells expressing *F9* variants derived from hemophilia B patients (Extended Data Table 1). Transduction with AAV vectors expressing SpCas9 (D10A) conjugated with BE4max or TadCBEd resulted in C>T conversion and a 2- to 3-fold increase in FIX activity (Fig. 3d). The efficacies were higher for TadCBEd than for BE4max (Fig. 3d and 3e). We therefore employed TadCBEd for further *in vivo* experiments because of their effectiveness and low bystander effects.

### Induction of FIX R338Q by base editing in vivo

To examine increases in FIX activity by base editing *in vivo*, we generated mice in which a human *F9* cDNA was knocked into the first mouse *F9* codon (Extended Data Fig. 3). We generated an AAV8 vector harboring SpCas9 (D10A) conjugated with TadCBEd driven by a HCRhAAT chimeric promoter (an enhancer element of the hepatic control region of the *APOE/C1* gene and the human anti-trypsin promoter) to efficiently and specifically express the target gene in the liver (Fig. 4a). Two AAV8 vectors were intravenously injected into adult *F9* knock-in mice, and FIX activity and antigen levels were examined after the injection. Plasma FIX activity but not antigen level, was increased 5–6 fold (Fig. 4b). Furthermore, G:C to A:T editing in the liver occurred at a rate of approximately 60% in these mice (Fig. 4c). The G:C to A:T editing was not observed in other organs (Fig. 4d). A marginal A to G conversion was detected at two 5′ upstream nucleotides, but it did not cause an amino acid change (Fig. 4e). The increase in plasma FIX activity by the introduction of R338Q was also observed in knock-in mice harboring hemophilia B variants (c.1138G>A, c.1300G>A, or c.1334C>T) (Fig. 4f,g). Moreover, the intraperitoneal administration of the vector into pups from knock-in mice efficiently induced C>T conversion, resulting in an increase in FIX activity (Extended Data Fig. 4).

**Fig. 4.**
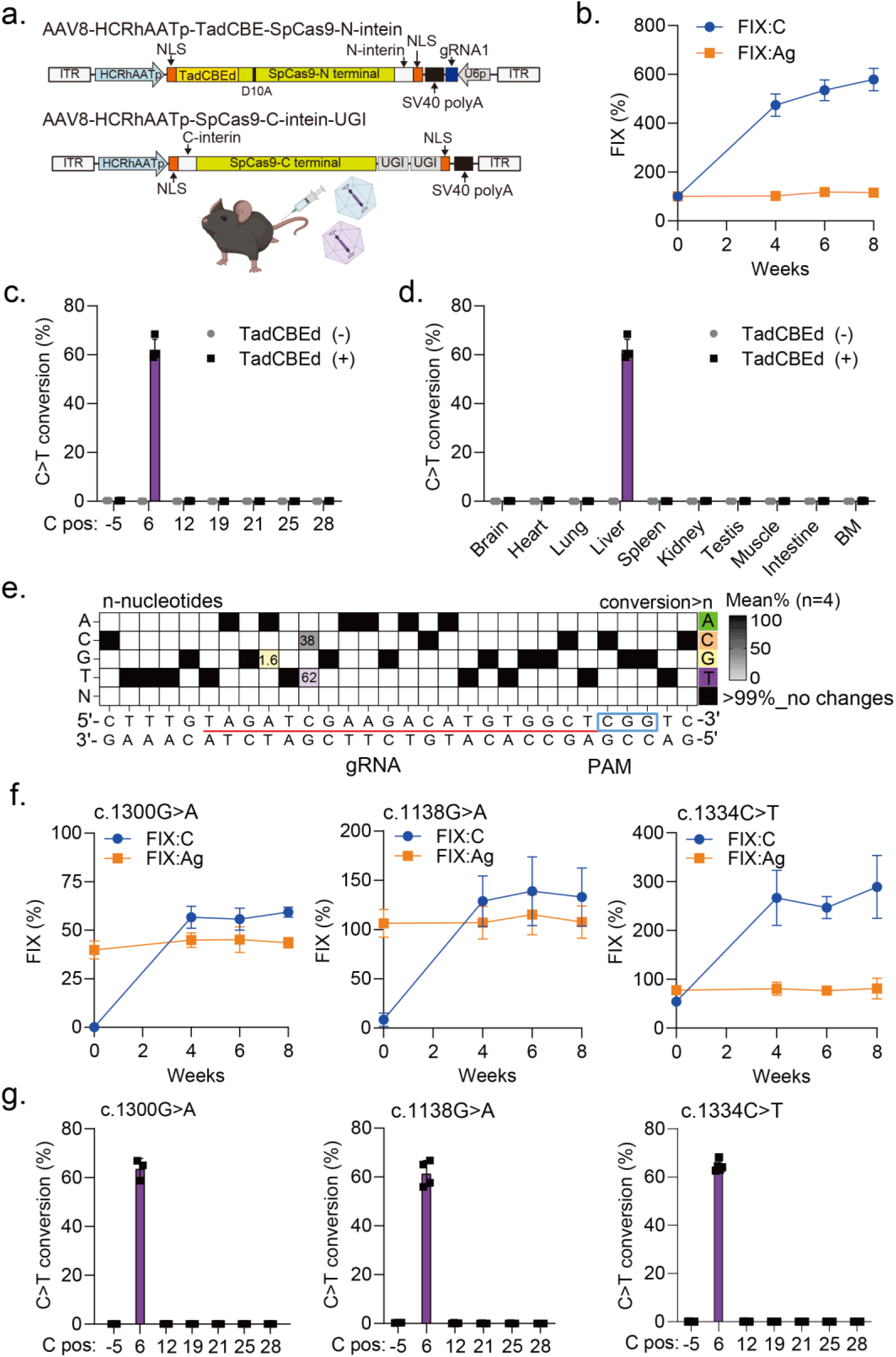
AAV vector mediated base editing *in vivo* increases FIX activity. Knock-in mice expressing a human *F9* cDNA were intravenously treated with two AAV8 vectors to express SpCas9 (D10A) conjugated with TadCBEd and UGI sequences, as well as gRNA1 sequence. **(a)** Schematic diagram of the AAV vector system. **(b)** Plasma FIX activity (FIX:C) and antigen (FIX:Ag) in adult knock-in mice after vector injection (1 × 10^12^ vg/body) [mean ± SD (n = 4)]. **(c)** Induction of C>T editing of DNA around the target site in liver assessed by NGS [mean ± SD (n = 4, treated group; n = 2, untreated group)]. Induction of C>T editing in DNA of the indicated organ [mean ± SD (n = 4)]. **(e)** The mean conversion frequency other than C>T substitution in liver DNA [mean ± SD (n = 4, treated group; n = 2, untreated group)]. **(f)** Plasma FIX:C and FIX:Ag in mice expressing *F9* cDNAs with hemophilia B variants after vector injection (1 × 10^12^ vg/body) [mean ± SD (n = 5–6)]. **(g)** Induction of C>T editing of DNA at the target site assessed by NGS [mean ± SD (n = 3–5)].

AAV vectors are an excellent platform for the delivery of target genes to the liver, but for genome editing, the expression of genome editing tools needs to be sustained. In addition, the induction of neutralizing antibodies against AAV vectors hampers re-administration of the same AAV vector formulation^24^. Therefore a particular AAV serotype can only be administered once. Indeed, all knock-in mice receiving the AAV vector for genome editing produced neutralizing antibodies against the AAV vector (Extended Data Fig. 5). We therefore employed lipid nanoparticles (LNPs) embedded with mRNA of the base editor to enable transient expression and re-administration for FIX R338Q introduction. *In vitro* transcribed mRNA is recognized by cells as foreign and triggers an immune response^25^. To avoid this immune response, cytosine (C) and uridine (U) in artificial mRNAs have been replaced with 5-methylcytosine (5MeC) and pseudouridine (*Ψ*U), respectively^26^. We compared the effects of such chemical modifications [5MeC, 5MeC+N1Metyl*Ψ*U, 5MeC+5-methoxy uridine (5MoU), *Ψ*U, N1Metyl*Ψ*U, and 5MoU] on the LNP delivery of *in vitro* transcribed mRNA into liver. As shown in Extended Data Fig. 6, the N1Metyl*Ψ*U modification gave the highest luciferase expression.

**Fig. 5.**
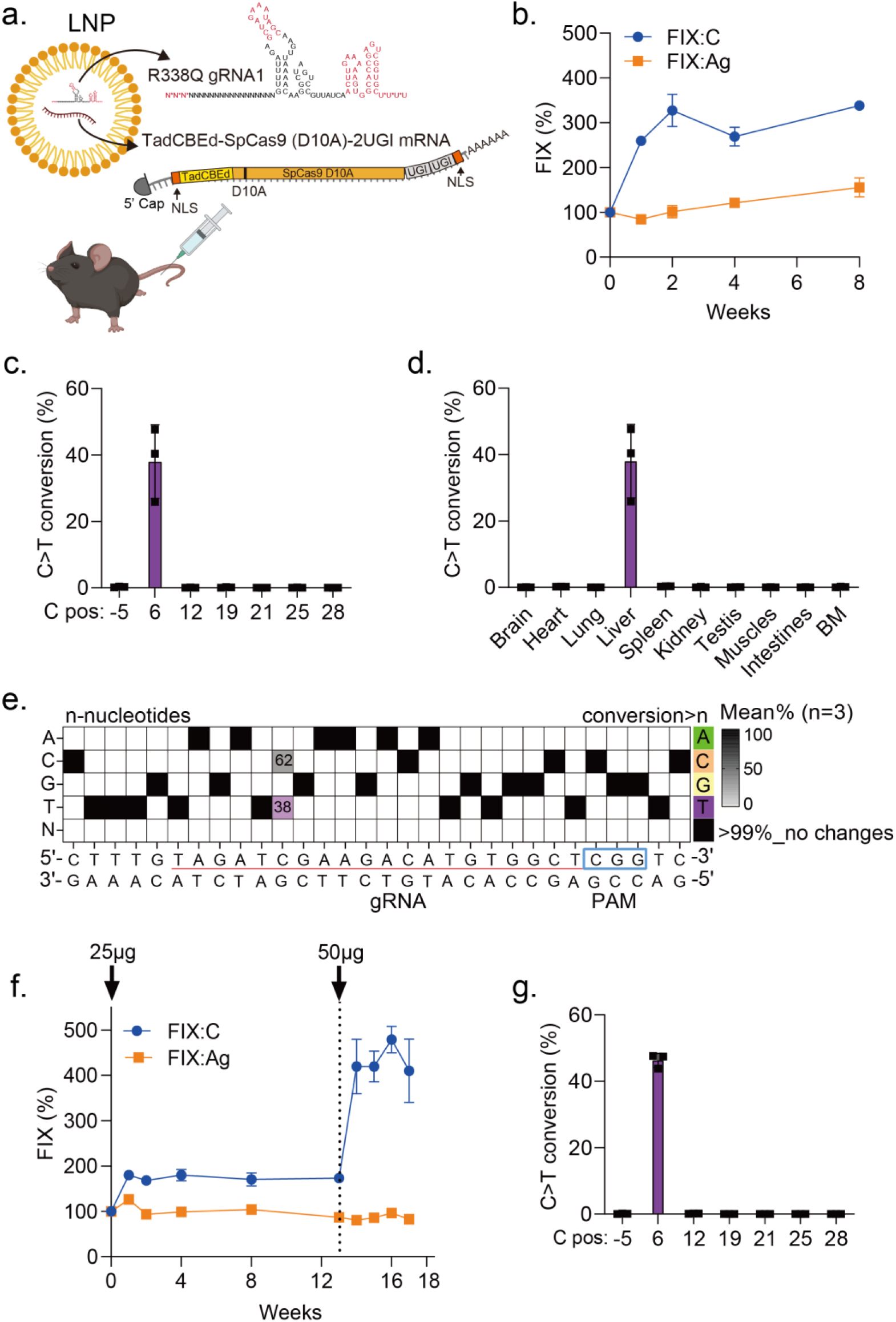
LNPs harboring base editor components increase FIX activity *in vivo*. Knock-in mice expressing a wild-type human *F9* cDNA were intravenously treated with LNPs harboring SpCas9 mRNA (D10A) conjugated with TadCBEd and UGI sequences, as well as gRNA1. **(a)** Schematic diagram of LNP injection. **(b)** Plasma FIX activity (FIX:C) and antigen (FIX:Ag) in adult knock-in mice after LNP injection (50 µg/body) [mean ± SD (n = 3)]. **(c)** Induction of C>T editing of DNA around the target site in liver assessed by NGS [mean ± SD (n = 3)]. **(d)** Induction of C>T editing in DNA of the indicated organ [mean ± SD (n = 3)]. **(e)** The mean conversion frequency other than C>T substitution in liver DNA. **(f, g)** Re-administration of LNPs. Knock-in mice expressing a wild-type human *F9* cDNA were first treated with LNPs (25 µg/body), and then with a higher dose of LNPs (50 µg/body) 13 weeks after the first injection. The changes in plasma FIX:C and FIX:Ag [mean ± SD (n = 3)]. **(g)** Induction of C>T editing of DNA at the target site assessed by NGS [mean ± SD (n = 3)].

We generated LNPs embedded with mRNAs of codon-optimized TadCBEd-SpCas9 with the N1Metyl*Ψ*U modification and the gRNA for the introduction of R338Q into *F9* (Fig. 5a). We intravenously administrated the LNPs into adult knock-in mice expressing human *F9* cDNA (Fig. 5a). As expected, plasma FIX activity, but not antigen level, increased significantly and persisted for at least 8 weeks after LNP injection (Fig. 5b). The efficiency of G:C to A:T base editing in *F9* mRNA was 30%– 40% in the liver (Fig. 5c). Base editing in other organs was marginal (Fig. 5d). Unexpected editing at target sites, including G:C to C:G or T:A to C:G, was not observed (Fig. 5e).

LNPs for mRNA delivery contain polyethylene glycol (PEG) for stabilization. PEG-specific antibodies can be induced by LNP treatment and may enhance clearance of PEG-containing drugs^27^. We therefore assessed the emergence of PEG-specific antibodies after LNP treatment to examine the possibility of re-administration. A very low level of anti-PEG IgM was transiently induced in mice treated with LNPs embedded with the base editor mRNA, but IgG against PEG was not observed (Extended Data Fig. 7), indicating that treatment with LNPs could be repeated. Indeed, a second administration of LNPs embedded with base editor mRNA further enhanced FIX activity (Fig. 5f,g).

## Discussion

Genome editing is gaining attention as next-generation treatment for genetic diseases. *Ex vivo* genome editing therapies targeting *BCL11A* for thalassemia and sickle cell disease were approved in 2023^28^. In addition, *in vivo* genome editing is currently under assessment in clinical trials for transthyretin amyloidosis, hereditary angioedema, and CEP290-related retinal degenerative diseases^3,4,29^. Base editing targeting a *PCSK9* splice donor site to reduce LDL cholesterol is also underway for hyperlipidemia^30^. All of these genome editing therapies aim to cause loss of gene function. However, base editing and prime editing to change individual amino acids, and knock-in or homologous recombination approaches to insert a gene into a target site are also important technologies for the therapeutic application of genome editing.

This study proposes a new concept of therapeutic genome editing, in which a gain-of-function variant is introduced by base editing. Most genetic diseases are caused by a single nucleotide variant; therefore, numerous inherited disorders are candidates for base editing. Applying prime editing would further expand the range of treatable patients. However, it is difficult to prepare personalized therapeutics for the variant present in each patient. We have introduced a gain-of-function variant in *F9* by G:C to A:T base editing *in vivo*, which improved the phenotype of a disease caused by a variety of variants. This concept of introducing a gain-of-function variant has the potential to be applied to the treatment of other genetic diseases. For example, K978C in CFTR, whose loss of function results in cystic fibrosis, increases protein function four-fold.^31^ To apply this approach to a wide range of diseases, it is important to find gain-of-function variants by structural analysis, artificial intelligence, and directed evolution.

Our strategy is a revolutionary concept that could provide a cure for hemophilia B patients. Hemophilia B gene therapy using AAV vectors is currently being developed, and some formulations have already been approved by the US Food and Drug Administration, The European Medicines Agency, and other authorities^7,32^. Current AAV vector gene therapy is a gene replacement therapy, in which a functional gene is expressed from its promoter sequence. AAV vectors reside in the nucleus as concatemers, and gene expression persists for at least several years in non-dividing cells. However, as cells proliferate, the vectors become diluted, resulting in a loss of efficacy^33^. Therefore, the effect of AAV gene therapy does not last long in childhood, when hepatocytes are proliferating^34^. Our strategy induces a single nucleotide variant in the genome; therefore, it is expected to have a lasting effect and to be able to treat the disease even in the neonatal period.

What proportion of the hemophilia B population would be eligible for this treatment? The prevalence of hemophilia B has been estimated at 3.8 cases in 100,0000 males^12^. Hemophilia can be divided into mild, moderate, and severe, based on FIX activity. Mild and moderate disease is more common (71%) than severe (FIX activity <1%)^12^. Our strategy can increase coagulation factor activity in patients with mild or moderate disease. Some patients with severe disease have low coagulation factor activity, but coagulation factor antigens are present; these are known as cross-reactive material-positive cases^35^. This corresponds to a molecular abnormality and is often caused by a single nucleotide variant^36^. Our strategy can be expected to have a therapeutic effect even in severe cases with cross-reactive material. Indeed, we demonstrated efficacy in knock-in mice in which FIX antigen was expressed but FIX activity was undetectable. This concept therefore has the potential to offer therapeutic benefit to more than 70% of hemophilia B patients, including mild, moderate, and severe with cross-reactive material-positive cases.

In the present study, we employed AAV vector and LNP delivery technologies. Both are used for the delivery of genome editing tools in human clinical trials. Although AAV vectors are a proven modality for gene delivery, there are several concerns over their use for genome editing targeting the liver. The most important drawback is the persistent expression of the antigenic bacterial Cas9 protein^37^. Furthermore, AAV vectors cannot be re-administered because neutralizing antibodies are generated after the first injection^38^. In contrast, genome editing with LNPs embedded with mRNAs and gRNAs allows transient expression of the genome editing tools and can be applied repeatedly. Our concept of universal base editing with LNPs allows for the initial administration of low doses of genome editing tools. This prevents coagulation factor activity being too high and causing thrombosis, and allows for increased efficacy by re-administration to achieve adequate coagulation factor activity. Furthermore, base editing with LNPs is advantageous in terms of genome toxicity. AAV vectors can integrate into genomic DNA^39^, the long-term safety of which is not known, but it can lead to clonal expansion of transduced hepatocytes in dogs treated with high-dose AAV vectors^39^. LNP-based genome editing avoids such vector-related genomic toxicity because the genome editing tools are transiently expressed. In addition, base editing avoids genome toxicity caused by large deletions and translocations at DSBs of target sites^40^.

The present study has several limitations. First, base editing may not always be able to induce gain-of-function mutations in other proteins. In this study, there was no bystander effect because there were no C nucleotides, other than the target nucleotide, in the editing space. Prime editing should be employed if the same base exists in the vicinity of the target nucleotide or if edits other than C>T or A>G are required. Second, actual off-target effects in humans have not been verified. Our Guide-seq analysis showed that C nucleotides were not present in off-target sites in HEK293 cells, nor were marginal gRNA-independent off-targets detected by R-loop assays. However, it is difficult to predict the real off-target effects of *in vivo* genome editing therapies, and careful safety monitoring is required in clinical trials. Finally, concerns about the safety of genome editing using LNPs should be carefully analyzed. A human clinical trial of base editing using LNPs targeting *PCKS9* (VERVE 101) was suspended because of adverse reactions, including liver damage^41^. A subsequent trial (VERVE102) was performed using GalNac-conjugated LNPs to enhance the introduction of LNPs into the liver^30,42^. It is essential to develop LNPs with improved organ tropism and to examine their safety in large animals including non-human primates.

In conclusion, we have demonstrated a novel concept for genome editing therapeutics that is based on engineering gain-of-function into a target protein. This approach has the potential to enable genome-edited treatment of many diseases with a single product. The development of base editing and prime editing technologies, accompanied by improvements in genome editing tool delivery to various organs *in vivo* may lead to cures for many genetic diseases. Although technological innovations in genome editing are expected to open up numerous therapeutic possibilities, the ethical issues of the technology always need to be discussed widely within society.

## Online Methods

### Plasmid construction

A human coagulation factor IX (FIX) cDNA was inserted into pBApo-EF1α (Takara Bio) for expression in HEK293 cells. Each variant was introduced by site-directed mutagenesis PCR. pX330 was used for the transient expression of SpCas9^43^. A sequence encoding SpCas9-NG was provided by Prof. Nureki (The University of Tokyo) and introduced into pX330. Base editor sequences and a uracil glycosylase inhibitor (UGI) sequence were synthesized by VectorBuilder or Twist Bioscience and introduced into pX330 containing a nickase version of SpCas9 (D10A). gRNA sequences were inserted into the BsaI site of pX330 after oligonucleotide annealing. In addition, gRNA sequences driven by the U6 promoter were inserted into pEX-A2-J2 (Eurofin Genomics) or pBluescript SK-(Stratagene). A SaCas9 cDNA was codon-optimized by GenScript, and then inserted into pcDNA3 (Thermo Fisher). The D10A and N580A mutations were introduced by site-directed mutagenesis to create dead SaCas9. The gRNA sequence driven by the U6 promoter was generated by Integrated DNA Technologies and inserted into pCDNA3 harboring dead SaCas9. For base editing using AAV vectors, we employed an intein-mediated split SpCas9 system^44^. Expression cassettes consisting of a promoter, SpCas9 (D10A)-N terminal, N-intein, SV40 polyA, and gRNA driven by the U6 promoter, or a promoter, C-intein, SpCas9-C terminal, two UGI sequences, and SV40 polyA, were introduced into pAAV.

### Cell culture and plasmid transfection

AAVpro 293T cells (Takara Bio) and HEK293 cells (JCRB Cell Bank #JCRB9068 293) were maintained in Dulbecco’s modified Eagle’s medium (FujiFilm Wako), supplemented with 10% fetal bovine serum (Thermo Fisher Scientific) and GlutaMAX (Thermo Fisher Scientific). Huh-7 cells (Riken BRC #RCB1366) were cultured in RPMI 1640 (FujiFilm Wako) supplemented with 10% fetal bovine serum and GlutaMAX. Transfection of HEK293 cells with indicated plasmids was conducted using Lipofectamine 3000 Reagent (Thermo Fisher Scientific) according to the manufacturer’s instructions. To generate stable expression clones, 400 μg/mL G418 (InvivoGen) was added to the culture medium after transfection. Vitamin K (5 µg/ml, menatetrenone, Eisai) was added to the culture medium 24 h before supernatants were collected for FIX measurement.

### Measurement of FIX activity and antigen

FIX activity (FIX:C) was measured with a one-stage clotting-time assay using an automated coagulation analyzer (CS-1600, Sysmex). FIX antigen (FIX:Ag) in the supernatant was measured with an ELISA kit (VisuLizet^™^ Factor IX Antigen Kit, Affinity Biologicals). To detect hFIX:Ag in mouse plasma, we employed an in-house ELISA. Microplates were coated with an anti-human FIX antibody (CEDARLANE). After blocking with 5% casein, diluted plasma samples were incubated for 1 h at 37°C. Antigen binding was detected with an anti-human FIX antibody conjugated with horseradish peroxidase (Affinity Biologicals) and ABTS microwell peroxidase substrate (Seracare)^15^. FIX levels in the supernatant are expressed as a % of that in normal human plasma. Changes in FIX levels in mice after treatment are expressed as a % of that in knock-in mice harboring a wild-type human *F9* cDNA.

### Computational structure modeling

Structural models of FVIII and activated FIXa were generated using AlphaFold3. The binding interactions between FVIII and FIXa variants (wild-type R338, Shanghai R338Q, and Padua R338L) were predicted with AlphaFold-Multimer^45^. Binding affinities between the A2 domain of FVIII and the protease domain of FIXa were quantified by HADDOCK 3.0^46^. Docking results were presented as HADDOCK scores and electrostatic energy values to assess the strength and stability of the FVIII-FIXa complexes. Protein structures were visualized and analyzed using PyMOL molecular visualization software (version 2.6, Schrödinger, LLC).

### T7 endonuclease assay and amplicon sequencing

Genomic DNA was extracted using SimplePrep reagent for DNA (Takara Bio) or a DNeasy Blood and Tissue Kit (Qiagen). Total RNA was isolated using an RNeasy Mini Kit (Qiagen). cDNA was synthesized using the PrimeScript real-time polymerase chain reaction (PCR) Kit (Takara Bio, Shiga, Japan). To detect DSBs at target sites, DNA fragments were amplified using ExTaq DNA polymerase (Takara Bio) and treated with T7 endonuclease (Nippon Gene). The DNA fragments were analyzed using a microchip electrophoresis system (MCE-202 MultiNA, Shimadzu). When indicated, PCR fragments were purified and size-selected using Sera-Mag Select beads (Cytiva). The purified amplicons were then barcoded using NEBNext Multiplex Oligos for Illumina (New England Biolab) and amplified using a KAPA HiFi HotStart ReadyMix PCR Kit (KAPA Biosystems). The target amplicons were subjected 300 bp paired-end sequencing on an Illumina NovaSeq 6000 system at the Research Institute for Microbial Diseases at Osaka University, Japan. The data were analyzed using CRISPResso2^47^.

### Guide-seq

Guide-seq analysis was performed as previously reported, with minor modification^48^. Briefly, HEK293 cells were transfected with double-strand oligonucleotide (dsODN, 100 pmol) and 500 ng of pX330 using Nucleofector 4-D (Lonza). Genomic DNA was isolated using a DNeasy Blood and Tissue Kit (Qiagen). We confirmed the insertion of dsODN in each experiment by NdeI restriction analysis. Genomic DNA was then fragmented using an M220 Focused-ultrasonicator (Covaris). Y-adaptor containing i5 (P5) was ligated using a GenNext NGS Library Prep Kit (Toyobo). Two PCR reactions were subsequently conducted to add an i7 (P7) sequence. The libraries were subjected 300 bp paired-end sequencing on Illumina MiSeq. The data were analyzed by GitHub code for Guide-seq. The primer and oligonucleotide for the analysis were identical to those used by Malinin et al^48^.

### AAV vector production

We employed AAV6 and AAV8 for *in vitro* and *in vivo* assays, respectively. The AAV genes were packaged by triple plasmid transfection of AAVpro293T cells (Takara Bio). AAV vectors were purified from the transfected cells by ultracentrifugation, as described previously^49^. The titer of AAV vectors was determined by quantitative PCR of the SV40 polyadenylation signal.

### Animal experiments

All animal experiments were approved by the Institutional Animal Care and Concern Committee of Jichi Medical University and were conducted in accordance with the committee’s guidelines (permission number 23067-01). Animal care was conducted according to the committee’s guidelines and the ARRIVE guidelines^50^.

C57BL/6 mice were purchased from SLC Japan (Shizuoka, Japan). Knock-in mice expressing human *F9* (*hF9*) with the R338L variant [C57BL/6-*F9*^*tm1*.*1(hF9_R338L)Tsuka*^, Riken BRC #RBRC12089] were developed by gene targeting of ES cells^15^. We generated several C57BL/6 mouse lines expressing human *hF9* cDNA by Cas9-mediated genome editing. First, we generated knock-in mice expressing a wild-type human *F9* (*hF9*) cDNA. A ribonucleoprotein complex consisting of SpCas9 protein (Alt-R S.p.HiFiCas9 Nuclease V3, Integrated DNA Technologies) and gRNA (Integrated DNA Technologies) together with a single-stranded oligonucleotide donor (Alt-R HDR Donor Oligo, Integrated DNA Technologies) was introduced into homozygous C57BL/6-*F9*^*tm1*.*1(hF9_R338L)Tsuka*^ embryos by electroporation (NEPA21 Type II, NepaGene). To create c.1138G>A, c.1300G>A, and c.1334C>T, homozygous C57BL/6-*F9*^*tm1*.*1(hF9_R338L)Tsuka*^ embryos were treated with an AAV6 donor vector for homologous recombination, and then the ribonucleoprotein complex was introduced by electroporation. The gRNA, single-stranded oligonucleotide donor, and AAV vector sequence with inverted terminal repeats are described in Extended Data Table 2.

Mice were maintained in isolators in the specific pathogen-free facility of Jichi Medical University at 23°C ± 3°C with a 12:12-h light/dark cycle. Mice were anesthetized with isoflurane (1%– 3%) for vector injection and to obtain blood samples, which were drawn from the jugular vein with a 29G micro-syringe (TERUMO, Tokyo, Japan) containing 1/10 (volume/volume) sodium citrate. Platelet-poor plasma was isolated by centrifugation and then frozen and stored at −80°C until analysis. AAV vectors or LNPs were administered intravenously to adult mice through the jugular vein. AAV vectors were administered intraperitoneally to neonatal mice.

### LNP production

Luciferase and SpCas9 (D10A) mRNAs conjugated with TadCBEd and two UGI sequences were synthesized by Elixirgen Scientific (Kanagawa, Japan). gRNA for SpCas9 targeting R338Q with chemical modifications^51^ was synthesized by Ajinomoto Bio-Pharma (Ibaraki, Japan). mRNA was loaded into LNPs using microfluidic mixing methods. An ethanol solution containing an ionizable lipid, 1,2-distearoyl-*sn*-glycero-3-phosphocholine (DSPC), cholesterol, and 1,2-dimyristoyl-*rac*-glycero-3-methoxypolyethylene glycol-2000 (PEG-DMG) at the molar ratio, 50:3:47:1.5, was prepared with a total lipid concentration of 8 mM. The RNA cargo [for base editing, 1:1 weight ratio of SpCas9 (D10A) mRNA:gRNA] was dissolved in 50 mM citrate buffer (pH 4.0), resulting in an RNA cargo concentration of 73.3 µg/mL. The lipid ethanol solution and the RNA solution were rapidly mixed using a glass-based iLiNP device^52^ at a total flow rate of 5 mL/min and an RNA-to-lipid flow rate of 3. The nitrogen-to-phosphate ratio was adjusted to 6. The resulting LNP solution was then dialyzed for 2 h or more at 4°C against 20 mM Tris HCl buffer (9% sucrose, pH 7.40) using Slide-A-Lyzer™ G3 Dialysis Cassettes (MWCO 20 kDa, Thermo Fisher Scientific). The LNP solution was concentrated by ultrafiltration using an Amicon Ultra-15 unit (MWCO 100 kDa, Millipore). The size and polydispersity of the LNPs were measured using a Zetasizer Nano ZS ZEN3600 instrument (Malvern Instruments, Worcestershire, UK). The encapsulation efficiency and total concentration of mRNA were measured by a Ribogreen assay, as described previously^53^. The LNPs were frozen and stored at −80°C until use.

### In vivo imaging

To analyze luciferase expression in the liver *in vivo*, mice were anesthetized with isoflurane. They then received the luciferin substrate (3 mg/body) intraperitoneally. Photons transmitted through the body were analyzed using an IVIS Imaging System and Living Image software (Xenogen, Alameda, CA). Quantitative data are expressed as photon units (photons/s).

### Measurement of anti-AAV neutralizing antibody

Plasma anti-AAV neutralizing antibody was assessed as described previously^54^. Briefly, plasma from animals was heat-inactivated (56°C, 30 min) and diluted with FBS, and then incubated with an AAV vector harboring luciferase at 37°C for 60 min. The mixture was then added to wells of a 96-well plate seeded with Huh-7 cells at multiplicity of infection of 1,500. The luciferase activity in cell lysate was determined at 48–72 h after the transduction. The samples were serially diluted with FBS, and then the inhibition of vector transduction was assessed. We estimated antibody titers that neutralized 50% vector transduction by nonlinear regression using GraphPad Prism (GraphPad, San Diego, CA).

### Measurement of anti-PEG antibody

Mouse serum anti-PEG IgG and anti-PEG IgM were measured using an Anti-PEG IgG ELISA kit (NTS-002-G, Nano T-sailing) and an Anti-PEG IgM ELISA kit (NTS-002-M, Nano T-sailing), respectively. Serum samples diluted 1:100 in Tris-buffered saline (TBS) containing 1% BSA were applied to wells of a 96-well plate coated with 1,2-distearoyl-sn-glycero-3-phosphoethanolamine-*n*-[methoxy (polyethylene glycol)-2000] (mPEG2000-DSPE) and incubated for 1 h. After five washes with TBS containing 0.05% CHAPS, 100 µL of horseradish peroxidase-conjugated goat anti-mouse IgG or IgM (1 µg/mL) in TBS containing 1% BSA were added and incubated for 1 h, followed by five washes with TBS containing 0.05% CHAPS. Horseradish peroxidase detection was initiated with 100 µL *o-*phenylenediamine (1 mg/mL), and the reaction was terminated with 100 µL 2 M H_2_SO_4_. Absorbance was measured at 490 nm using a microplate reader (Sunrise, TECAN Japan). Serum anti-PEG IgG or IgM concentrations were quantified based on standard curves obtained using monoclonal mouse anti-PEG IgG (Clone HIK-G11) and anti-PEG IgM (Clone HIK-M11).

### Statistics

The number of independent biological samples or mice used in each experiment is specified in the corresponding figure legends. Data are presented as the mean ± SD. A two-tailed unpaired Student’s *t-* test was used for two-group comparisons. For experiments involving more than two groups, one-way ANOVA was performed, followed by Tukey’s *post hoc* test, depending on the experimental design. All statistical analyses were conducted using GraphPad Prism version 10.3.1 (GraphPad Software, San Diego, CA).

## Data availability

Source data underlying the Figures are provided as a Supplementary Data file. The data obtained from the current study are available from the corresponding author on reasonable request.

## Acknowledgements

The study was supported by grants JP24bm1223004 (T.O.), JP23am0401005 (O.N., T.O.), JP23fk0410037 (T.O.), JP24fk0410061 (T.O.), and JP24bm1323001 (T.O.) from the Japan Agency for Medical Research and Development (AMED), and a research grant from The Naito Foundation. We thank Hiromi Ozaki, Yuiko Ogiwara, Sachiyo Kamimura, Yaeko Suto, Mika Kishimoto, Tamaki Aoki, Mai Hayashi, Tomoko Noguchi, and Hiroko Hayakawa (Jichi Medical University) for their technical assistance. We acknowledge Dr. Daisuke Motooka in the NGS core facility at the Research Institute for Microbial Diseases of Osaka University for NGS sequencing and advice on data analysis. Some figures were created with Biorender.com. We also thank Jeremy Allen, PhD, from Edanz (https://jp.edanz.com/ac) for editing a draft of this manuscript.

## Author contributions

N.B. and Y.K.: the acquisition, analysis and interpretation of data, drafting the paper.

T.H.: the acquisition and interpretation of data.

R.I., R.S., Y.N., H.N., H.T., and TS: the acquisition of data.

M.H., K.B., T.T.: the analysis and interpretation of data.

Y.S. and T.I.: the acquisition and interpretation of data, drafting the paper

A.H. and O.N.: the acquisition and interpretation of data, revising the paper.

T.O.: study conception, the acquisition, analysis and interpretation of data, drafting and revising the paper.

All authors approved the submitted version and agreed to be personally accountable for their own contributions and for the accuracy and integrity of any part of the work.

## References

1. Wang, J.Y. & Doudna, J.A. CRISPR technology: A decade of genome editing is only the beginning. Science 379, eadd8643 (2023).

2. Zheng, Y., et al. Precise genome-editing in human diseases: mechanisms, strategies and applications. Signal Transduction and Targeted Therapy 9, 47 (2024).

3. Gillmore, J.D., et al. CRISPR-Cas9 In Vivo Gene Editing for Transthyretin Amyloidosis. New England Journal of Medicine 385, 493–502 (2021).

4. Longhurst, H.J., et al. CRISPR-Cas9 In Vivo Gene Editing of <i>KLKB1</i> for Hereditary Angioedema. New England Journal of Medicine 390, 432–441 (2024).

5. Papapetropoulos, A., et al. Novel drugs approved by the EMA, the FDA, and the MHRA in 2023: A year in review. British Journal of Pharmacology 181, 1553–1575 (2024).

6. Adikusuma, F., et al. Large deletions induced by Cas9 cleavage. Nature 560, E8–E9 (2018).

7. O’Leary, K. Base editors in the clinic. Nature Medicine 29, 2972–2972 (2023).

8. Landrum, M.J., et al. ClinVar: public archive of interpretations of clinically relevant variants. Nucleic Acids Res 44, D862–868 (2016).

9. Rees, H.A. & Liu, D.R. Base editing: precision chemistry on the genome and transcriptome of living cells. Nature Reviews Genetics 19, 770–788 (2018).

10. Newby, G.A., et al. Base editing of haematopoietic stem cells rescues sickle cell disease in mice. Nature 595, 295–302 (2021).

11. Musunuru, K., et al. In vivo CRISPR base editing of PCSK9 durably lowers cholesterol in primates. Nature 593, 429–434 (2021).

12. Iorio, A., et al. Establishing the Prevalence and Prevalence at Birth of Hemophilia in Males. Annals of Internal Medicine 171, 540–546 (2019).

13. Hemophilia, W.F.o. WFH Annual Global Survey 2021. (Montreal, Canada, 2021).

14. Johnsen, J.M., et al. Novel approach to genetic analysis and results in 3000 hemophilia patients enrolled in the My Life, Our Future initiative. Blood Advances 1, 824–834 (2017).

15. Hiramoto, T., et al. PAM-flexible Cas9-mediated base editing of a hemophilia B mutation in induced pluripotent stem cells. Communications Medicine 3, 56 (2023).

16. Simioni, P., et al. X-Linked Thrombophilia with a Mutant Factor IX (Factor IX Padua). New England Journal of Medicine 361, 1671–1675 (2009).

17. Samelson-Jones, B.J., et al. Evolutionary insights into coagulation factor IX Padua and other high-specific-activity variants. Blood Advances 5, 1324–1332 (2021).

18. Wu, W., et al. Factor IX alteration p.Arg338Gln (FIX Shanghai) potentiates FIX clotting activity and causes thrombosis. Haematologica 106, 264–268 (2021).

19. Samelson-Jones, B.J., Finn, J.D., George, L.A., Camire, R.M. & Arruda, V.R. Hyperactivity of factor IX Padua (R338L) depends on factor VIIIa cofactor activity. JCI Insight 4(2019).

20. Nishimasu, H., et al. Engineered CRISPR-Cas9 nuclease with expanded targeting space. Science 361, 1259–1262 (2018).

21. Chen, L., et al. Re-engineering the adenine deaminase TadA-8e for efficient and specific CRISPR-based cytosine base editing. Nature Biotechnology 41, 663–672 (2023).

22. Huang, J., et al. Discovery of deaminase functions by structure-based protein clustering. Cell 186, 3182-3195.e3114 (2023).

23. Doman, J.L., Raguram, A., Newby, G.A. & Liu, D.R. Evaluation and minimization of Cas9-independent off-target DNA editing by cytosine base editors. Nature Biotechnology 38, 620–628 (2020).

24. Baatartsogt, N., et al. Successful liver transduction by re-administration of different adeno-associated virus vector serotypes in mice. The Journal of Gene Medicine 25, e3505 (2023).

25. Chen, Y.G., et al. Sensing Self and Foreign Circular RNAs by Intron Identity. Molecular Cell 67, 228-238.e225 (2017).

26. Vaidyanathan, S., et al. Uridine Depletion and Chemical Modification Increase Cas9 mRNA Activity and Reduce Immunogenicity without HPLC Purification. Mol Ther Nucleic Acids 12, 530–542 (2018).

27. Chen, B.-M.Cheng, T.-L. & Roffler, S.R. Polyethylene Glycol Immunogenicity: Theoretical, Clinical, and Practical Aspects of Anti-Polyethylene Glycol Antibodies. ACS Nano 15, 14022–14048 (2021).

28. FDA Approves First Gene Therapies to Treat Patients with Sickle Cell Disease. (Food and Drug Administration (FDA), 2023).

29. Pierce Eric, A., et al. Gene Editing for CEP290-Associated Retinal Degeneration. New England Journal of Medicine 390, 1972–1984 (2024).

30. Kasiewicz, L.N., et al. GalNAc-Lipid nanoparticles enable non-LDLR dependent hepatic delivery of a CRISPR base editing therapy. Nature Communications 14, 2776 (2023).

31. Woodall, M., et al. Expression of gain-of-function CFTR in cystic fibrosis airway cells restores epithelial function better than wild-type or codon-optimized CFTR. Molecular Therapy Methods & Clinical Development 30, 593–605 (2023).

32. Wang, J.-H., Gessler, D.J., Zhan, W., Gallagher, T.L. & Gao, G. Adeno-associated virus as a delivery vector for gene therapy of human diseases. Signal Transduction and Targeted Therapy 9, 78 (2024).

33. Verdera, H.C., Kuranda, K. & Mingozzi, F. AAV Vector Immunogenicity in Humans: A Long Journey to Successful Gene Transfer. Molecular Therapy 28, 723–746 (2020).

34. Muhuri, M., Levy, D.I., Schulz, M., McCarty, D. & Gao, G. Durability of transgene expression after rAAV gene therapy. Molecular Therapy 30, 1364–1380 (2022).

35. W Keith Hoots, M.H. Genetics of hemophilia A and B. In: UpToDate. (Wolters Kluwer, 2023).

36. Castaman, G. & Matino, D. Hemophilia A and B: molecular and clinical similarities and differences. Haematologica 104, 1702–1709 (2019).

37. Wagner, D.L., et al. High prevalence of Streptococcus pyogenes Cas9-reactive T cells within the adult human population. Nature Medicine 25, 242–248 (2019).

38. George, L.A., et al. Long-Term Follow-Up of the First in Human Intravascular Delivery of AAV for Gene Transfer: AAV2-hFIX16 for Severe Hemophilia B. Molecular Therapy 28, 2073–2082 (2020).

39. Greig, J.A., et al. Integrated vector genomes may contribute to long-term expression in primate liver after AAV administration. Nature Biotechnology 42, 1232–1242 (2024).

40. Komor, A.C., Kim, Y.B., Packer, M.S., Zuris, J.A. & Liu, D.R. Programmable editing of a target base in genomic DNA without double-stranded DNA cleavage. Nature 533, 420–424 (2016).

41. Philippidis, A. StockWatch: Investors Less Forgiving than Analysts as Verve Tumbles. GEN Edge 6, 323–329 (2024).

42. Luo, Y., Hou, Y., Zhao, W. & Yang, B. Recent progress in gene therapy for familial hypercholesterolemia treatment. iScience 27(2024).

43. Cong, L., et al. Multiplex Genome Engineering Using CRISPR/Cas Systems. Science 339, 819–823 (2013).

44. Truong, D.J., et al. Development of an intein-mediated split-Cas9 system for gene therapy. Nucleic Acids Res 43, 6450–6458 (2015).

45. Abramson, J., et al. Accurate structure prediction of biomolecular interactions with AlphaFold 3. Nature 630, 493–500 (2024).

46. Giulini, M., et al. Towards the accurate modelling of antibody-antigen complexes from sequence using machine learning and information-driven docking. Bioinformatics 40(2024).

47. Clement, K., et al. CRISPResso2 provides accurate and rapid genome editing sequence analysis. Nature Biotechnology 37, 224–226 (2019).

48. Malinin, N.L., et al. Defining genome-wide CRISPR–Cas genome-editing nuclease activity with GUIDE-seq. Nature Protocols 16, 5592–5615 (2021).

49. Kashiwakura, Y., et al. Efficient gene transduction in pigs and macaques with the engineered AAV vector AAV.GT5 for hemophilia B gene therapy. Molecular Therapy Methods & Clinical Development 30, 502–514 (2023).

50. Percie du Sert, N., et al. The ARRIVE guidelines 2.0: Updated guidelines for reporting animal research. PLOS Biology 18, e3000410 (2020).

51. Finn, J.D., et al. A Single Administration of CRISPR/Cas9 Lipid Nanoparticles Achieves Robust and Persistent <em>Ina0;Vivo</em> Genome Editing. Cell Reports 22, 2227–2235 (2018).

52. Maeki, M., et al. Mass production system for RNA-loaded lipid nanoparticles using piling up microfluidic devices. Applied Materials Today 31, 101754 (2023).

53. Hashiba, K., et al. Overcoming thermostability challenges in mRNA–lipid nanoparticle systems with piperidine-based ionizable lipids. Communications Biology 7, 556 (2024).

54. Kashiwakura, Y., et al. The seroprevalence of neutralizing antibodies against the adeno-associated virus capsids in Japanese hemophiliacs. Molecular Therapy Methods & Clinical Development 27, 404–414 (2022).

