## Extended file 1 for "Universal base editing for hemophilia B"

### SUPPLEMENTAL MATERIALS

Extended Figure 1 A structural binding model of activated FIX and FVIII.
Extended Figure 2 R-loop assay to detect gRNA-independent off-target editing.
Extended Figure 3 Generation of human *F9* cDNA knock-in mice.
Extended Figure 4 Increase in FIX activity by AAV vector harboring a base editor in neonatal mice.
Extended Figure 5 Induction of anti-AAV neutralizing antibody in response to the AAV vector harboring a base editor.
Extended Figure 6 Assessment of mRNA modification *in vivo*.
Extended Figure 7 Induction of anti-PEG antibody in response to LNPs harboring a base editor.

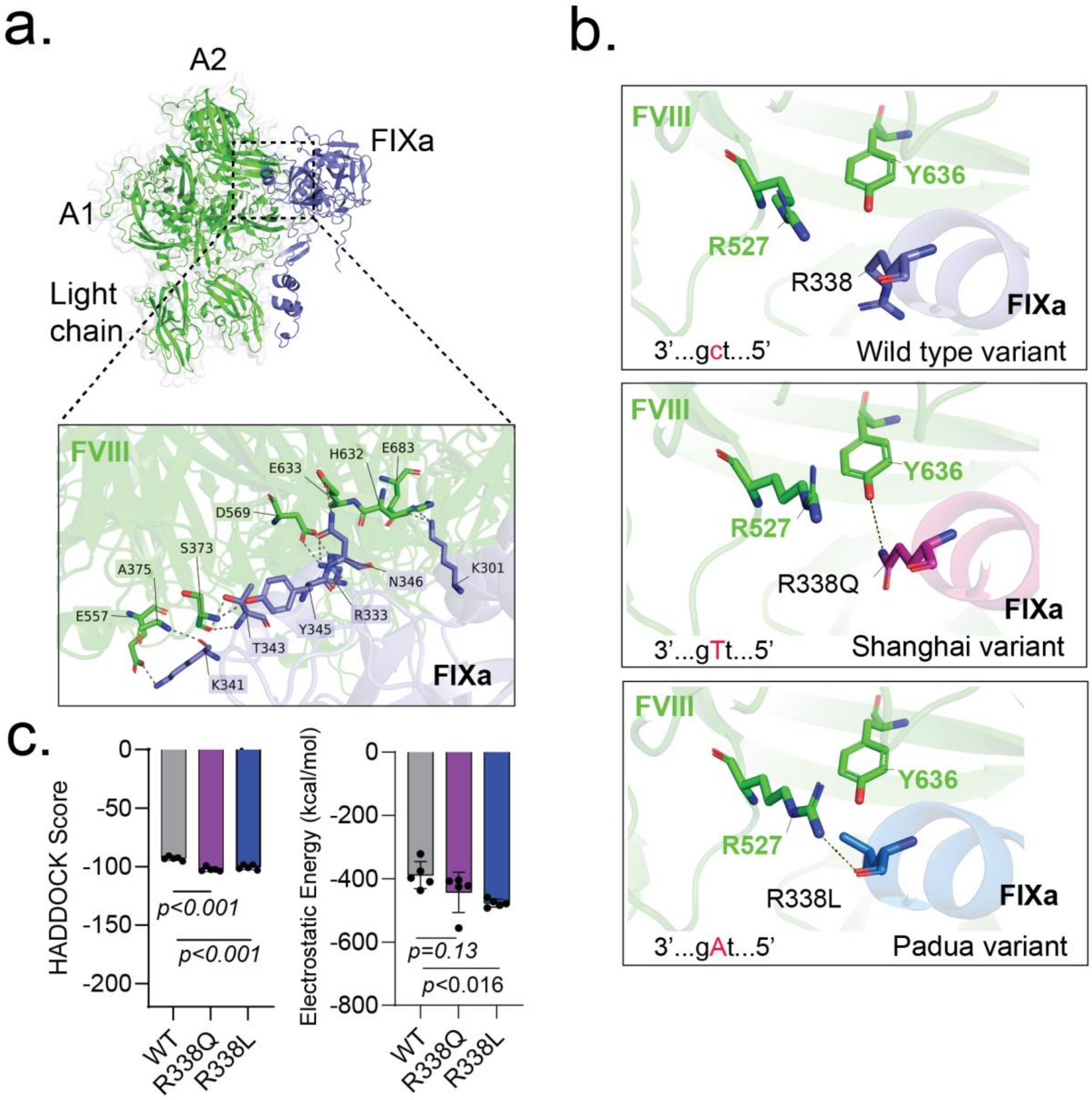


**Extended Figure 1. A structural binding model of activated FIX and FVIII. (a)** A structural binding model between activated FIX (FIXa) and FVIII assessed by AlphaFold 3 (green: FVIII; blue: FIXa). Magnified box shows the binding site and key residues involved in the binding between FIXa and the A2 domain of FVIII. **(b)** Close-up views of the R338 site in the wild type, the Shanghai variant (R338Q), and the Padua variant (R338L) of FIXa, illustrating specific interactions with FVIII residues. **(c)** Binding affinity scores and electrostatic energy values for FIXa-FVIII interaction [mean ± SD (n = 5)]. Statistical significance was assessed using one-way ANOVA followed by Tukey's *post hoc* test for pairwise comparisons.

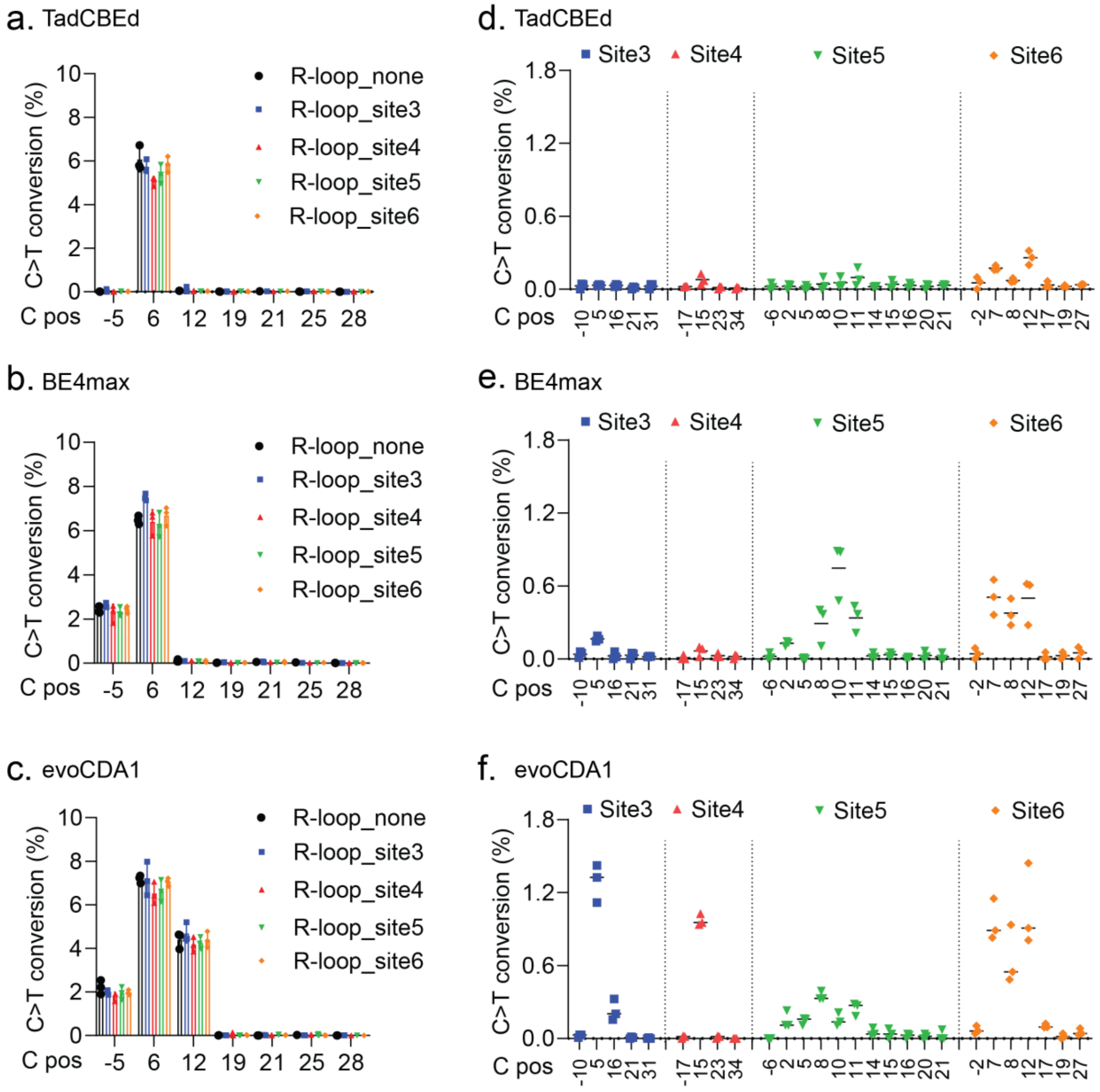


**Extended Figure 2 R-loop assay to detect gRNA-independent off-target editing.** HEK293 cells were transfected with the plasmid expressing SpCas9 (D10A) conjugated with a base editor (TadCBEd, BE4max, or evoCDA1) and gRNA1 for induction of R338Q, as well as the plasmid harboring dead SaCas9 (D10A and H840A) and gRNA sequence (site 3, 4, 5, or 6). **(a, b, c)** Induction of C>T editing of mRNA around the on-target site assessed by NGS [mean ± SD (n = 3)]. **(d, e, f)** The frequency of gRNA-independent off-target editing at sites 3, 4, 5, and 6 [mean ± SD (n = 3)].

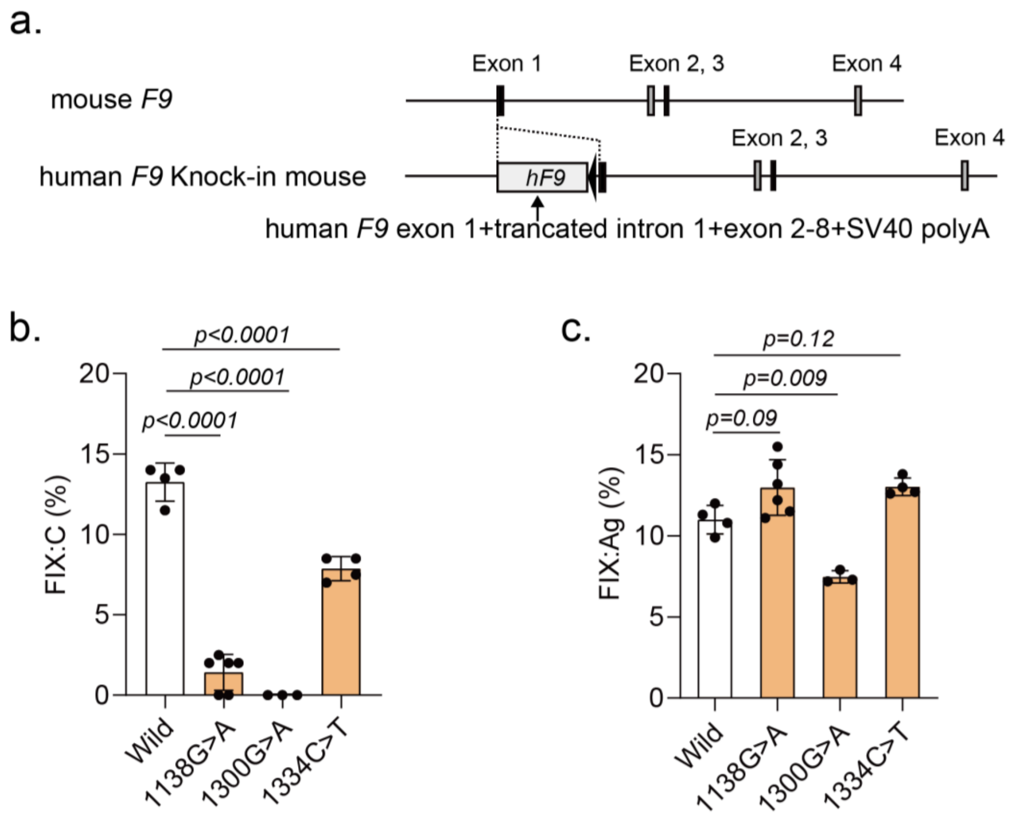


**Extended Figure 3 Generation of human *F9* cDNA knock-in mice. (a)** Schematic diagram of *F9* knock-in mice. **(b, c)** FIX activity (FIX:C) **(b)** and FIX antigen (FIX:Ag) **(c)** in knock-in mice [mean ± SD (n = 3–6)]. Statistical analysis was performed using one-way ANOVA followed by Tukey's *post hoc* test for pairwise comparisons.

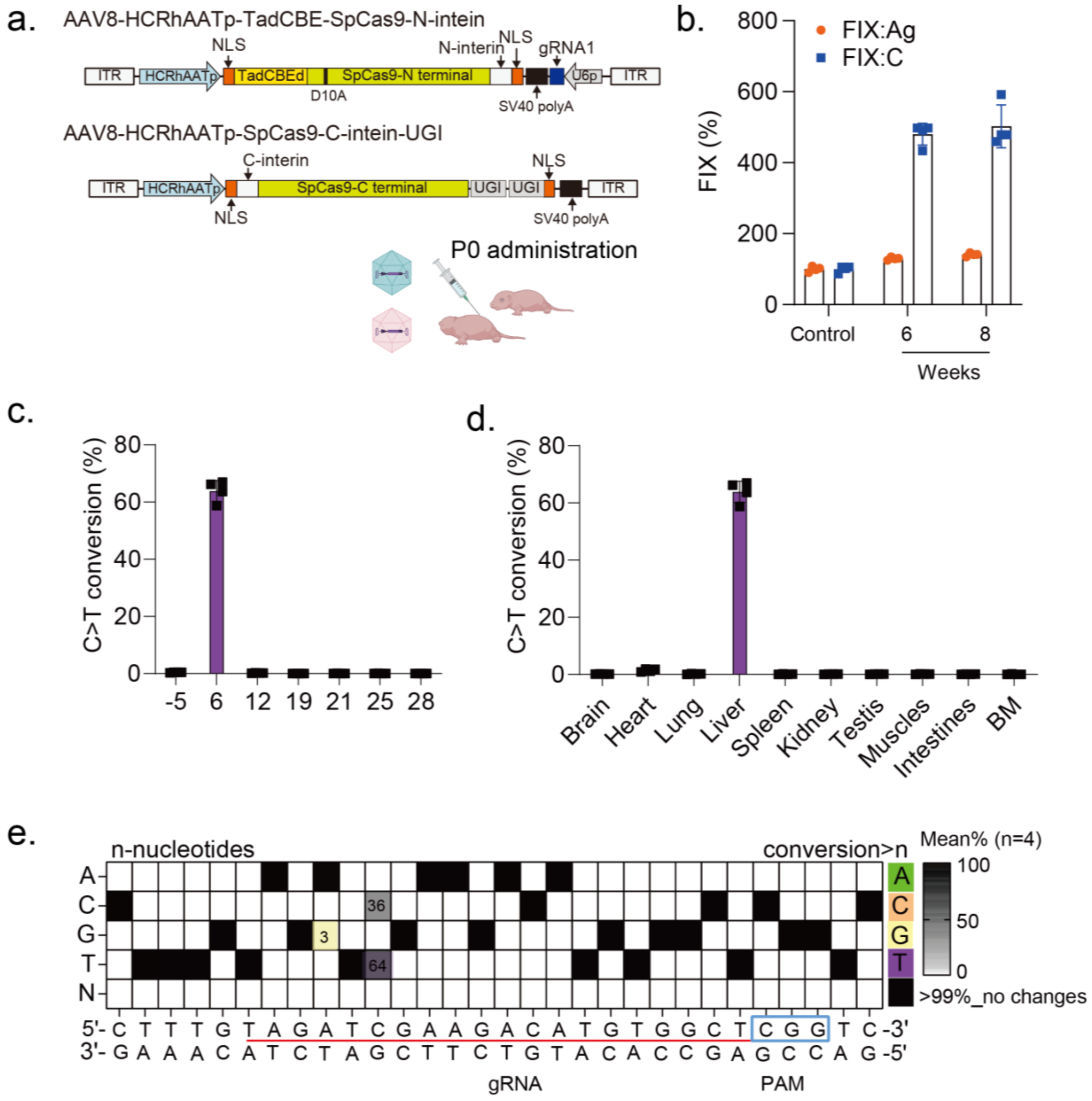


**Extended Figure 4 Increase in FIX activity by AAV vector harboring a base editor in neonatal mice.** Knock-in neonatal mice expressing a human *F9* cDNA were intraperitoneally treated with two AAV8 vectors to express SpCas9 (D10A) conjugated with TadCBEd and UGI sequences, as well as gRNA1 sequence. **(a)** Schematic diagram of the AAV vector system. **(b)** Plasma FIX activity (FIX:C) and antigen (FIX:Ag) after vector injection ($3 \times 10^{11}$ vg/body) [mean ± SD (n = 4)]. **(c)** Induction of C>T editing of DNA around the target site in liver assessed by NGS [mean ± SD (n = 4)]. **(d)** Induction of C>T editing in DNA of indicated organs [mean ± SD (n = 4)]. (**e**) The mean conversion frequency other than C>T substitution in liver DNA.

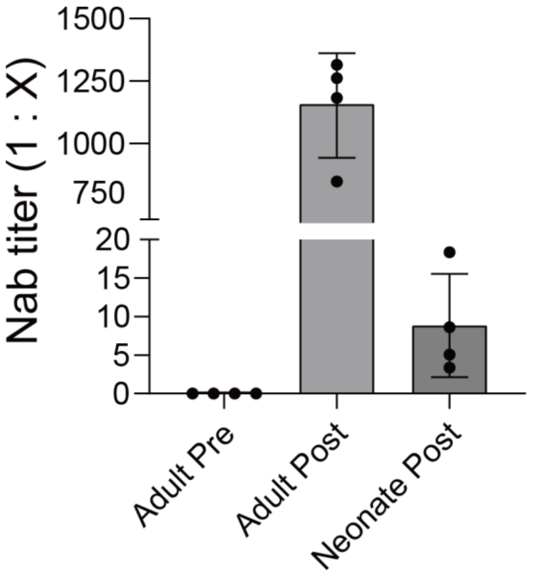


**Extended Figure 5 Induction of anti-AAV neutralizing antibody in response to the AAV vector harboring a base editor.** Knock-in mice (adults and neonates) expressing a human *F9* cDNA were treated with two AAV8 vectors to express SpCas9 (D10A) conjugated with TadCBEd and UGI sequences, as well as gRNA1 sequence. The plasma levels of anti-AAV8 neutralizing antibody (Nab) were determined by the ability to neutralize AAV8 vector transduction of Huh-7 cells [mean ± SD (n =4)].

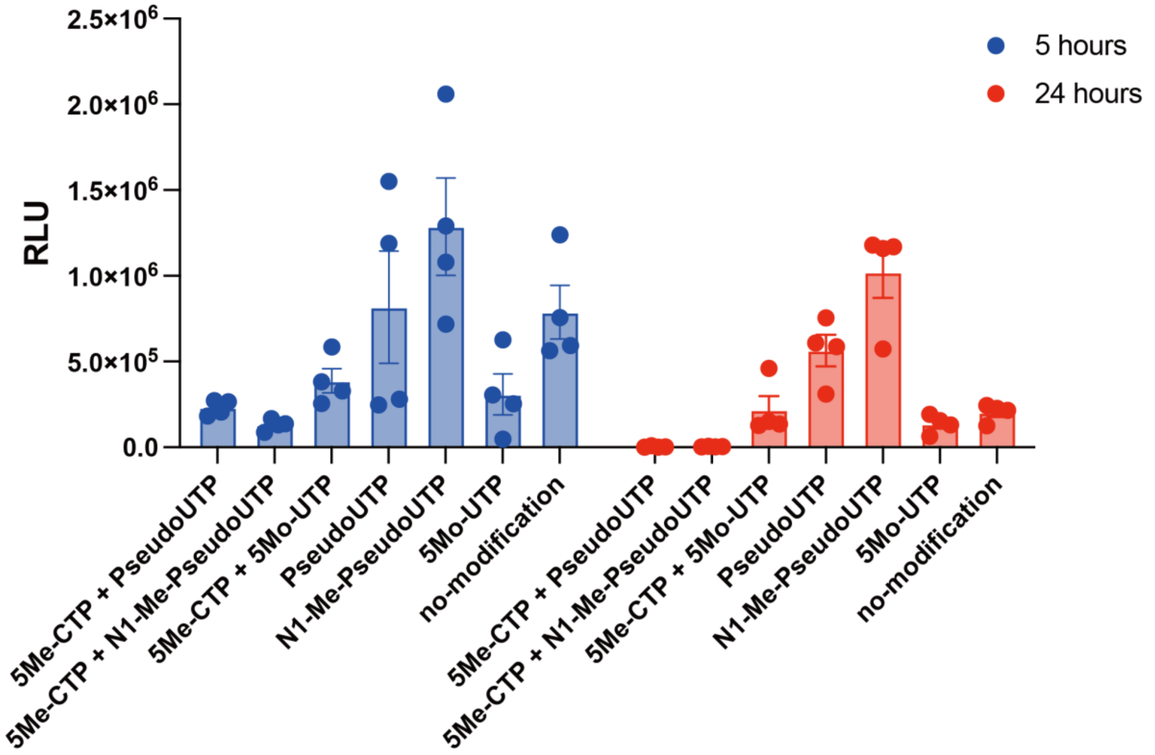


**Extended Figure 6 Assessment of mRNA modification *in vivo*.** Seven-week-old male C57BL/6 mice were treated with LNPs harboring luciferase mRNA (0.5 µg/body). Luciferase expression in the liver was assessed by *in vivo* imaging at 5 and 24 h after LNP injection. Comparison of *in vivo* luciferase expression among mRNAs with several modifications [mean ± SD (n = 4)].

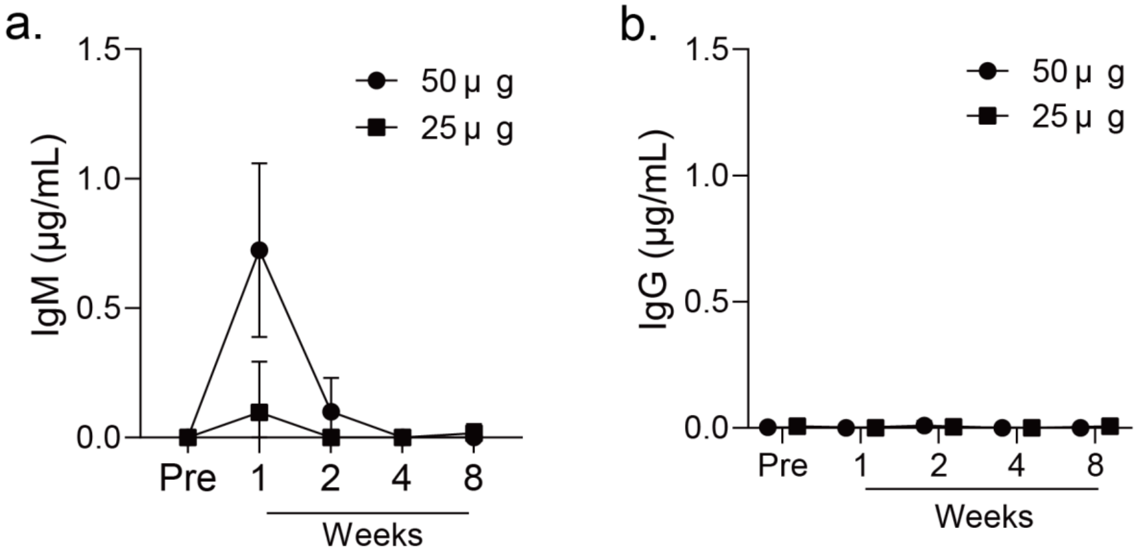


**Extended Figure 7 Induction of anti-PEG antibody in response to LNPs harboring a base editor.** Knock-in mice expressing a wild-type human *F9* cDNA were intravenously treated with LNPs harboring mRNA of SpCas9 (D10A) conjugated with TadCBEd and UGI sequences, as well as gRNA1 (25 or 50 µg/body). Plasma levels of anti-PEG IgM **(a)** and IgG **(b)** were measured over time by ELISA [mean ± SD (n = 4)].
